# Muscle fiber proteomics reveals sex- and fiber type-specific adaptations to resistance training

**DOI:** 10.1101/2024.09.16.612737

**Authors:** Lukas Moesgaard, Roger Moreno-Justicia, Johann Schmalbruch, Søren Jessen, Ben Stocks, Jens Bangsbo, Atul S. Deshmukh, Morten Hostrup

**Affiliations:** The August Krogh Section for Human Physiology, Department of Nutrition, Exercise and Sports, University of Copenhagen, Denmark; Novo Nordisk Center for Basic Metabolic Research, University of Copenhagen, Denmark; Department of Molecular Medicine and Surgery, Integrative Physiology, Karolinska Institutet, Stockholm, Sweden

## Abstract

Skeletal muscle hypertrophy is a hallmark of resistance training that positively impacts health and longevity. However, despite physiological differences between sexes and fiber types, the underlying proteome changes with resistance training have not been studied in a sex- and fiber type-specific manner. Herein, we show sex differences in the fiber type-specific proteome, predominantly in type II fibers. Following 8 weeks of resistance training, substantial remodeling of the human skeletal muscle proteome occurred in a sex- and fiber type-specific manner. Notably, type II fibers exhibited much greater adaptations across both sexes, whereas the main sex-difference was a greater remodeling of intermediate filaments in females. In addition, baseline abundance of proteins involved in translation was highly correlated with fiber hypertrophy, and differed between sexes and fiber types. Thus, translational capacity may partially explain differences in resistance training-induced hypertrophy. Our findings demonstrate key aspects of sex- and fiber type differences in muscle physiology and their contributions to resistance training-induced adaptions.

## Introduction

Skeletal muscle function is essential for health and longevity. Greater muscle strength is associated with a reduced risk of all-cause mortality, with this association even stronger in females than males^1^. Resistance exercise training is a key intervention for maintaining health and longevity, via increased muscle mass and strength, as well as improved metabolic health^2-4^. However, the underlying molecular remodeling in response to resistance training remains incompletely described.

Heterogeneous in nature, skeletal muscle is composed of both type I fibers, which have slow contractile properties and are more oxidative, and type II fibers, which have fast contractile properties and are more glycolytic^5^. Notably, this mixture of fiber types affects the response to resistance training, as greater muscle hypertrophy occurs in type II fibers than type I fibers^6-8^. Furthermore, we recently demonstrated sex-specific resistance training-induced hypertrophy of type I fibers^9^. Thus, our understanding of skeletal muscle adaptations to training can greatly benefit from research in both sexes. Unfortunately, the number of studies in biological research that only include male participants outnumber the number of studies that include females by a large margin^10^.

Studying skeletal muscle adaptations in a fiber type-specific manner is challenging, due to the need to isolate and identify specific fiber types. Studies that have investigated the fiber type-specific response to resistance training have used targeted approaches, such as immunohistochemistry^11^ or western blotting^12^ to investigate single proteins of interest. Naturally, this must be seen as a limitation as muscle adaptations are complex, and single proteins are unlikely to be representative of the changes occurring in the entire muscle proteome. By quantifying several thousands of proteins in a single run, the application of mass spectrometry-based proteomics to study resistance training can disentangle complex molecular adaptations with unprecedented resolution. While a few studies have investigated changes in the human skeletal muscle proteome following resistance training^13-15^, it remains unknown whether remodeling of the human skeletal muscle proteome is sex- and fiber type-specific.

In this study, we applied our recently developed muscle fiber proteomics workflow^16^ to investigate sex- and fiber type-specific effects of resistance training. Our analysis revealed that proteome remodeling occurred primarily in type II fibers, and that intermediate filament protein abundance primarily increased in females. Furthermore, we discovered differences between sexes and fiber types in the abundance of proteins involved in translation, which may have implications for resistance training-induced hypertrophy. These findings demonstrate key differences in male and female skeletal muscle, and underline the importance of sex-specific research in the field of physiology.

## Results

### Study overview and proteomics workflow

Twelve females and 12 males who did not engage in resistance training prior to the intervention completed 8 weeks of supervised resistance training performed 3 times a week, with muscle biopsies sampled from *m. vastus lateralis* before and 3 days after the last training bout (Fig. 1A). The resistance training intervention increased leg lean mass (*p* < 0.001; Fig. 1B) and strength, assessed by 1-repetition maximum (1RM) leg press (*p* < 0.001; Fig. 1C), in both males and females. After freeze-drying the muscle biopsies, single fibers were dissected and typified using dot blotting with antibodies against MYH2 (type II) and MYH7 (type I). Fibers were then pooled by participant and time point, resulting in pools of 16 ± 6 (mean ± SD) type I fibers and 21 ± 6 type II fibers that were prepared for proteomics analysis.

**Fig. 1.**
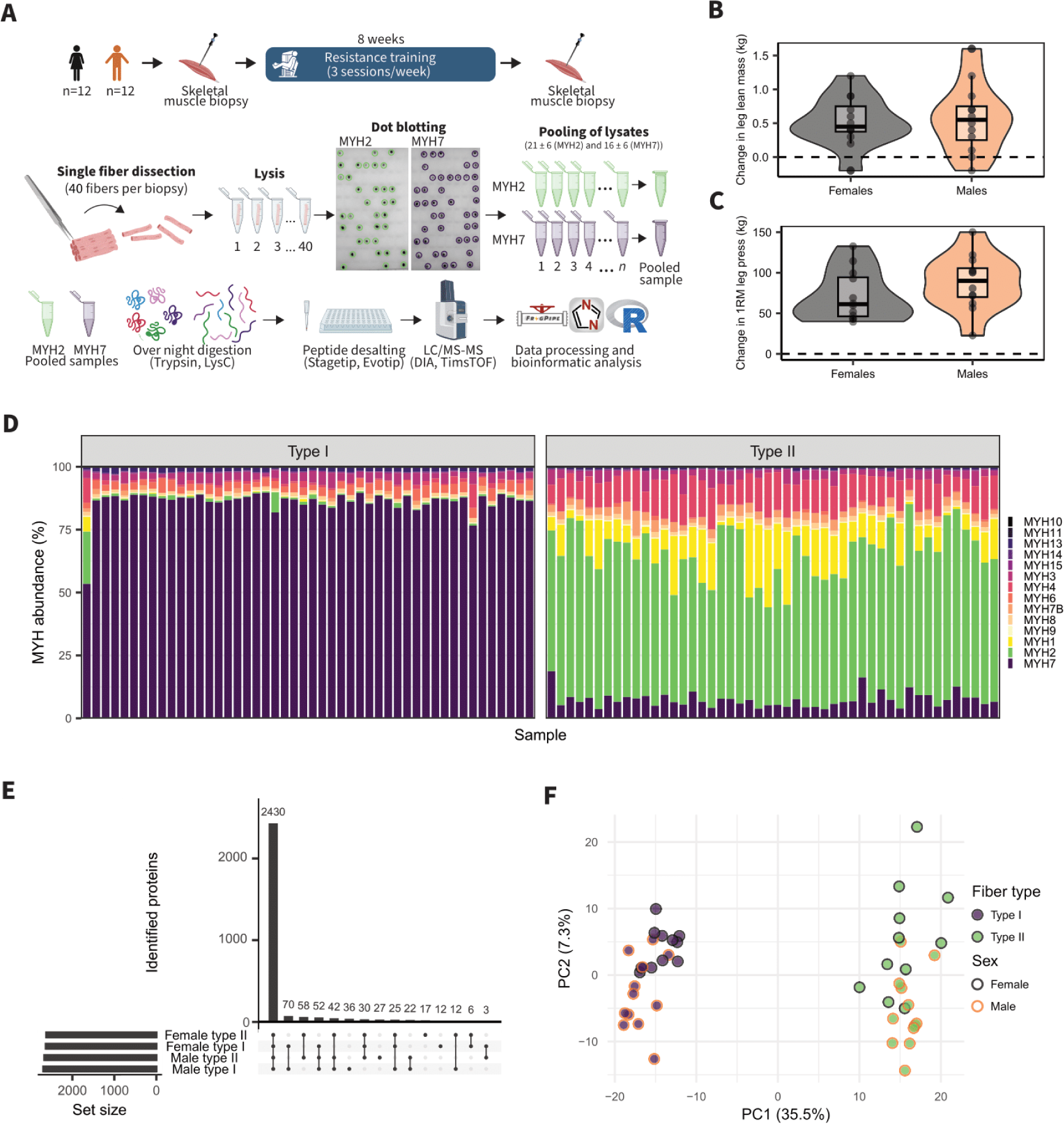
Overview of study design, proteomics workflow and data validation. **(A)** Study overview. **(B)** Change in leg lean mass following 8 weeks of resistance training. **(C)** Change in 1-repetition maximum (1RM) following 8 weeks of resistance training. **(D)** Abundance of MYH isoforms in type I and type II fibers pools. **(E)** Upset plot of identified proteins. **(F)** Principal component analysis (PCA) of baseline samples.

For preparation for proteomics analysis, samples were digested and peptides were measured by liquid chromatography tandem mass-spectrometry (LC-MS/MS) using timsTOF DIA-PASEF. This approach detected a total of 3197 proteins, of which 2883 proteins were identified in at least 70% of samples in either male or female type I or type II fiber pools, respectively, and used for bioinformatics analysis (Table S1). Proteomics validated the purity of type I and type II fiber pools, as characterized by high abundance of MYH7 in type I fibers and high abundance of MYH2 in type II fibers (Fig. 1D). While the majority of the proteome was quantified in each sex and fiber type, 22 proteins were unique to male fibers and 6 proteins were unique to female fibers. In addition, 70 proteins were unique to type I fibers and 58 proteins were unique to type II fibers (Fig. 1E). In addition, principal component analysis of baseline samples revealed that fiber types clearly separated along principal component 1, which accounted for 35.5% of total variance between samples, whereas sexes seemed to separate along principal component 2, which accounted for 7.3% of the variance between samples (Fig. 1F). This identifies fiber type and sex as the main determinants of variance in human skeletal muscle.

### Fiber type differences in the skeletal muscle proteome

To dive deeper into the differences in the sex- and fiber type-specific proteome, we compared only baseline samples to avoid the confounding effects of resistance training. As expected, myosin heavy chains, myosin light chains, and troponin isoforms were the main drivers of separation between type I and type II fibers within principal component 1 (Fig. 2A). When directly comparing fiber types, we identified 1004 proteins that were differentially expressed between type I and type II fibers in females and 1656 proteins that were differentially expressed in males (FDR < 0.05; Fig. 2B; Table S2). There was a large overlap in fiber type-specific proteins between males and females, although the difference between fiber types was quantitatively greater in males (Fig. 2C). In both sexes, type I fibers were enriched for gene ontology biological process (GO:BP) terms related to mitochondrial ATP synthesis, while type II fibers were enriched for NADH regeneration, driven by the glycolytic enzymes ENO2, ENO3, PFKL, PFKM, PGAM2, PGK1, PGK2, PKM, and TPI1. However, other terms such as mitochondrial gene expression and cytoplasmic translation were only enriched in male type I and type II fibers, respectively (FDR < 0.05; Fig. 2D). Thus, the difference between fiber types appears moderately more distinct in males than females.

**Fig. 2.**
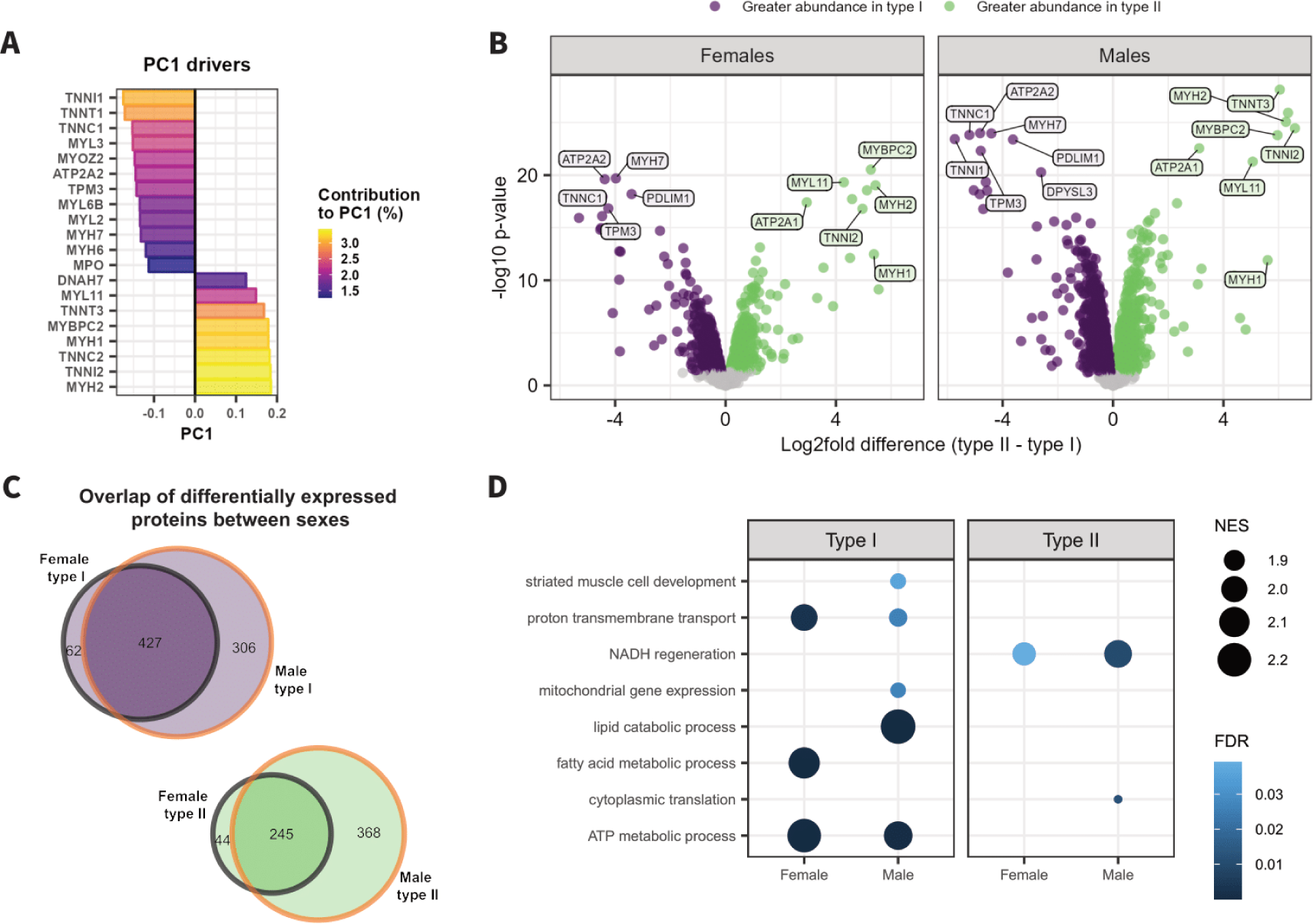
Sex-specific differences between fiber types. **(A)** Principal component 1 drivers and their contributions to variance between baseline samples. **(B)** Volcano plot showing differential expression in protein abundance between type I and type II fibers in males and females. Differentially expressed proteins (FDR < 0.05) are marked with purple (higher abundance in type I fibers) and green (higher abundance in type II fibers). **(C)** Euler diagram of the overlap of differentially expressed proteins between males and females. **(D)** Dot plot of gene set enrichment analysis showing normalized enrichment score (NES) of enriched gene ontology biological process (GO:BP) terms in type I and type II fibers in males and females (FDR < 0.05).

### Proteomics reveals sex-differences in the fiber type-specific proteome

Exploring the differences between male and female fibers, separation along principal component 2 was mainly driven by keratins and other intermediate filament proteins (Fig. 3A), which in skeletal muscle form a complex cytoskeletal network that connects Z-disks to nuclei^17,18^ and costameres at the sarcolemma^19,20^. Importantly, these differences were mainly driven by differences in type II fibers as only 8 proteins were differentially expressed between sexes in type I fibers, whereas 141 proteins were differentially expressed between sexes in type II fibers (FDR < 0.05; Fig. 3B; Table S3–4).

**Fig. 3.**
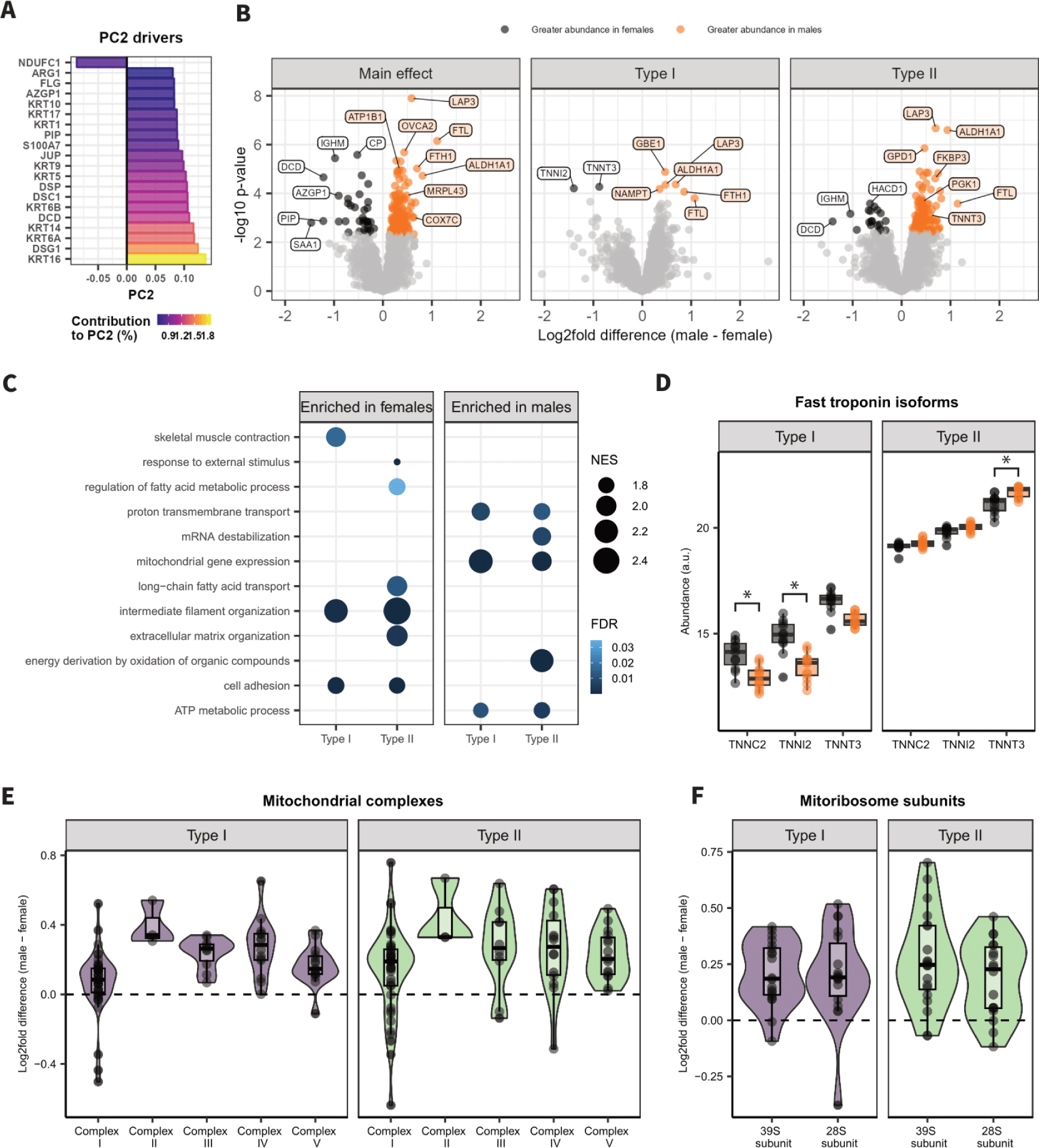
Fiber type-specific proteome differences between males and females. **(A)** Principal component 2 drivers and their contributions to variance between baseline samples. **(B)** Volcano plot showing differential expression in protein abundance between males and females. First displayed as a main effect, then specifically in type I and type II fibers. Differentially expressed proteins (FDR < 0.05) are marked with black (higher abundance in females) and orange (higher abundance in males). **(C)** Dot plot of gene set enrichment analysis showing normalized enrichment score (NES) of enriched gene ontology biological process (GO:BP) terms in males and females in type I and type II fibers (FDR < 0.05). **(D)** Boxplot of fast troponin isoforms in males and females in type I and type II fibers. *FDR < 0.05. **(E)** Log2fold difference between males and females of proteins in mitochondrial complexes. **(F)** Log2fold difference between males and females of proteins in mitoribosome subunits. Each circle represents a protein.

Gene set enrichment analysis confirmed that female skeletal muscle had a greater abundance of intermediate filament proteins irrespective of fiber type, whereas male fibers were enriched for GO:BP terms related to mitochondria (FDR < 0.05; Fig. 3C). Some GO:BP terms were also enriched in a both sex- and fiber type-specific manner. This included terms such as mRNA destabilization that was specifically enriched in male type II fibers, as well as extracellular matrix organization and GO:BP terms related to fatty acid metabolism that were specifically enriched in female type II fibers (FDR < 0.05; Fig. 3C). Interestingly, female type I fibers were enriched for the term skeletal muscle contraction. This term was driven by differences in fast troponin isoforms, where abundance of TNNI2 and TNNT3 in type I fibers was greater in females, but abundance of TNNT3 in type II fibers was greater in males (FDR < 0.05; Fig. 3D). This suggests that females have faster contractile properties of type I fibers, but slower type II fibers, indicating that contractile properties of fiber types are also more distinct in males than females.

Given that terms related to mitochondria were enriched in male skeletal muscle, irrespective of fiber type, we used Mitocarta 3.0 ^21^ to assess whether this was specific to certain mitochondrial proteins or whether it applies to overall greater mitochondrial density. Protein abundance tended to be higher in males than females for most proteins in all mitochondrial complexes, although the difference was more variable in complex I (Fig. 3E). Likewise, the abundance of proteins within both mitoribosome subunits was higher in males than females (Fig. 3F). Taken together, these data indicate that males possess a higher skeletal muscle oxidative capacity than females.

### Remodeling of the muscle proteome following resistance training is sex- and fiber type-specific

We next assessed whether resistance training would elicit a sex- and fiber type-specific response. Here, we observed remarkable remodeling of the skeletal muscle proteome with many of these changes being specific to sex and fiber type. Overall, resistance training upregulated 313 proteins and downregulated 139 proteins (main effect: FDR < 0.05; Fig. 4A; Table S5). However, when assessed in a sex- and fiber type-specific manner only 9 and 7 proteins were regulated in female and male type I fibers, respectively. In contrast, 164 proteins were regulated in female type II fibers and 101 proteins were regulated in male type II fibers (FDR < 0.05; Fig. 4B; Table S6). Interestingly, these proteomics data mirror greater resistance training-induced hypertrophy in type II fibers^6-8^.

**Fig. 4.**
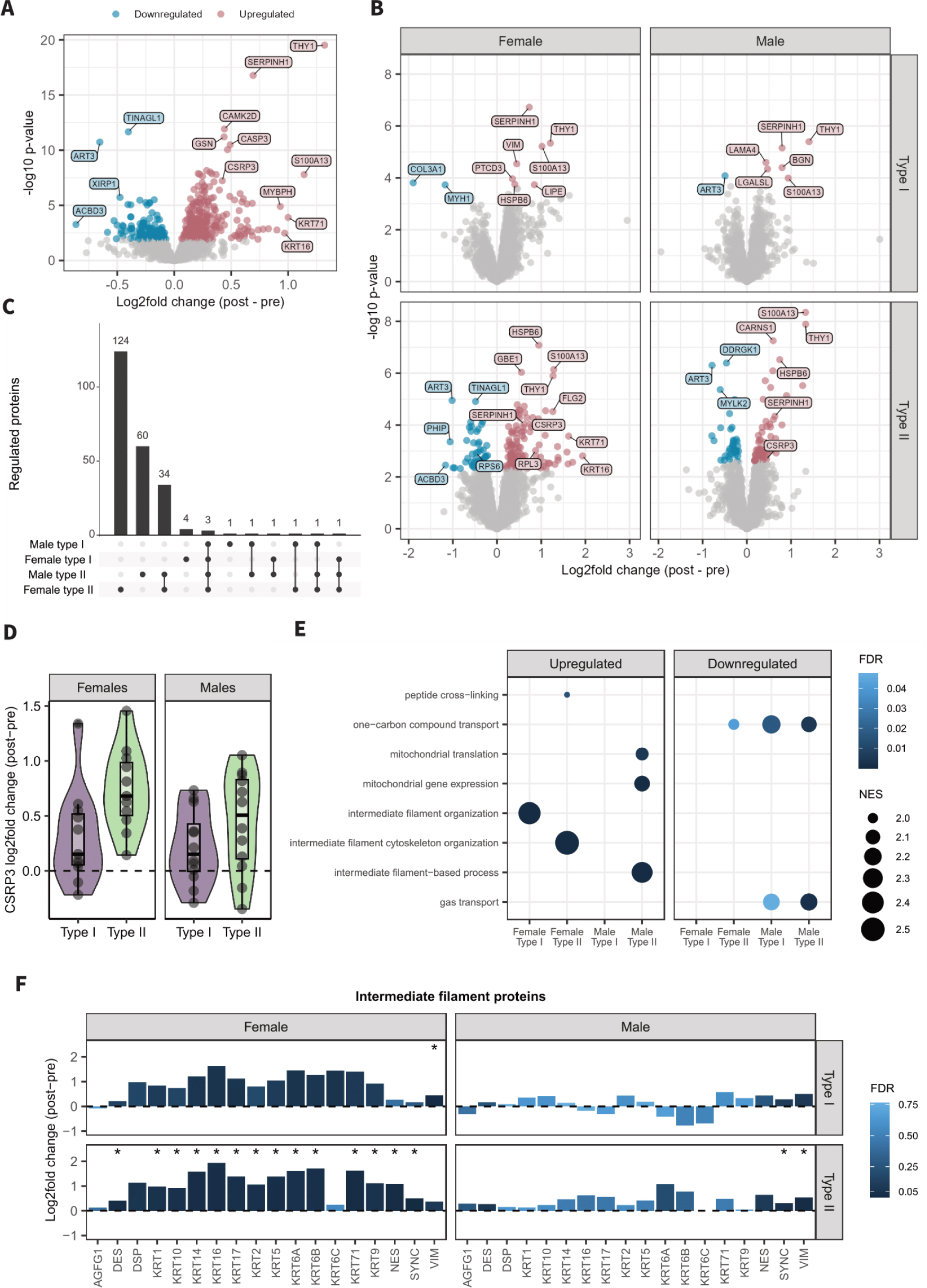
Sex- and fiber type-specific changes in the proteome following 8 weeks of resistance training. **(A)** Volcano plot showing a main effect of resistance training on protein abundance. **(B)** Volcano plot showing sex- and fiber type-specific changes in protein abundance following resistance training. Regulated proteins (FDR < 0.05) are marked with blue (downregulated) and red (upregulated). **(C)** Upset plot of regulated proteins. **(D)** Plot of individual changes in abundance of CSRP3 in male and female type I and type II fibers. **(E)** Dot plot of gene set enrichment analysis showing normalized enrichment score (NES) of upregulated and downregulated gene ontology biological process (GO:BP) terms in male and female type I and type II fibers (FDR < 0.05). **(F)** Plot of log2fold change in abundance of intermediate filament proteins in male and female type I and type II fibers. *FDR < 0.05.

To further explore adaptations specific to type II fibers, we investigated the 34 proteins that were regulated in both male and female type II fibers, but not type I fibers (Fig. 4C). Here, we identified upregulation of CSRP3 (Fig. 4D), which is associated with the GO:BP term “skeletal muscle tissue development”. In addition, CSRP3 is associated with the GO:BP term “detection of muscle stretch”, which implies a role as a mechanosensor regulating skeletal muscle hypertrophy^22^.

To elucidate the observed sex- and fiber type-specific remodeling of the skeletal muscle proteome we used gene set enrichment analysis, which revealed that the GO:BP terms mitochondrial translation and mitochondrial gene expression were upregulated in male type II fibers only (FDR < 0.05; Fig. 4E). Terms related to intermediate filaments were upregulated in both female type I and type II fibers, but only male type II fibers (FDR < 0.05; Fig. 4E). Furthermore, intermediate filament proteins were upregulated to a substantially larger degree in female skeletal muscle, irrespective of fiber type (Fig. 4F). Given that the largest differences in the baseline proteome were also related to intermediate filaments and mitochondria, these data indicate that resistance training induces divergence in the sex-specific proteome.

### Baseline protein content regulating translation correlates with fiber hypertrophy and is sex- and fiber type-specific

To further explore differences in the skeletal muscle proteome underlying differential hypertrophy we assessed correlations between protein abundance at baseline and fiber hypertrophy assessed as the change in fiber cross-sectional area (CSA) by immunohistochemistry^9^ (Fig. 5A). Here, gene set enrichment analysis revealed that the strongest correlated proteins with fiber hypertrophy were enriched for the GO:BP term translation (FDR < 0.05, Fig. 5B–C). This term was primarily driven by ribosomal proteins, eukaryotic initiation factors, and eukaryotic elongation factors, but also included kinases involved in translation such as mTOR, RPS6KA3, and RPS6KB2. Thus, fibers with greater translational capacity seem to experience greater resistance training-induced hypertrophy.

**Fig. 5.**
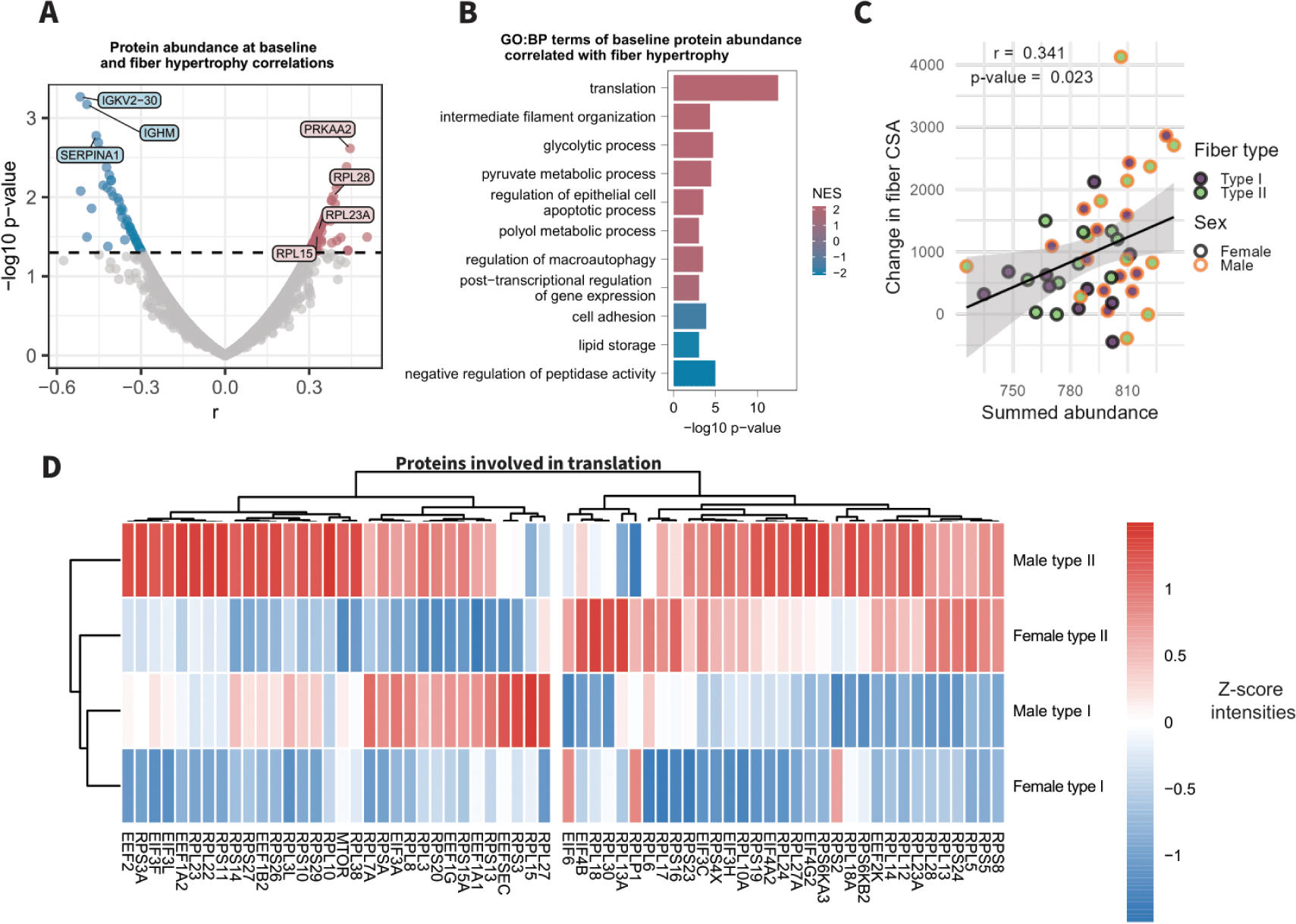
Differences in the proteome that correlate with muscle fiber hypertrophy following 8 weeks of resistance training. **(A)** Pearson’s correlation coefficient between abundance of individual proteins at baseline and the change in fiber cross-sectional area (CSA) ^9^. **(B)** Plot of gene set enrichment analysis showing normalized enrichment score (NES) and –log10 p-value of gene ontology biological process (GO:BP) terms at baseline correlated with the change in fiber CSA (FDR < 0.05). Positive correlations are marked with red and negative correlations are marked with blue. **(C)** Scatter plot showing Pearson’s correlation between summed abundance of proteins involved in translation and the change in fiber CSA^9^. **(D)** Heatmap and hierarchical clustering of proteins involved in translation. Red indicates high abundance and blue indicates low abundance.

As translational capacity was the strongest predictor of fiber hypertrophy, we performed hierarchical clustering of proteins involved in translation, which revealed two clusters. One cluster was driven by fiber type, where abundance was higher in both male and female type II fibers than type I fibers, whereas the other cluster was driven by sex, where abundance was higher in male fibers than female fibers (Fig. 5D). As such, these differences in translational capacity may explain previously observed sex- and fiber type-specific resistance training-induced hypertrophy^9^.

## Discussion

It is widely recognized that resistance training is the gold standard intervention for inducing muscle hypertrophy^23^, which is of utmost importance for physical function and metabolic health^24^. However, despite obvious physiological differences between males and females, as well as the consistent observation of fiber type-specific hypertrophy following resistance training^6-8^, the underlying sex- and fiber type-specific remodeling of the proteome has yet to be mapped. Herein, we demonstrate substantial fiber type-specific differences between males and females in the skeletal muscle proteome and its remodeling following resistance training. Furthermore, we build on these findings by elucidating differences in the skeletal muscle proteome that likely explain resistance training-induced sex- and fiber type-specific hypertrophy.

While we have previously shown differences in the proteome between type I and type II fibers in males^16^, our results show that the skeletal muscle proteome is both sex- and fiber type-specific. Notably, our results indicate that female type I and type II fibers are less distinct than male fiber types. This applied both to contractile properties that were driven by sex differences in fast troponin isoforms, as well as biological processes such as mitochondrial metabolism and cytosolic translation. Our findings also suggest greater oxidative capacity of skeletal muscle in males than females, irrespective of fiber type. This is consistent with other physiological differences between sexes such as greater blood volume and hemoglobin levels in males^25,26^, that ultimately result in a higher maximal oxygen uptake. In contrast, greater abundance of intermediate filament proteins likely play a greater role in providing cellular integrity^27^ in female skeletal muscle, suggesting structural differences in skeletal muscle between sexes.

Resistance training resulted in substantial remodeling of the skeletal muscle proteome that was highly fiber type-specific. Here, our findings display that greater hypertrophy of type II fibers^6-8^ following resistance training is reflected by type II muscle fiber-specific remodeling of the proteome. CSRP3 was one such protein exclusively upregulated in type II fibers. CSRP3 is a mechanosensitive protein involved in myogenesis and satellite cell-mediated myonuclear accretion^28^. Upon activation, CSRP3 translocates from the z-disk of sarcomeres to nuclei and interacts with muscle transcription factors (MFRs) such as MyoD, MRF4, and myogenein^29-31^. Accordingly, absence of nuclear CSRP3 blunts mechanical strain-induced protein accretion in cultured neonatal rat myocytes^29^, and knockdown of CSRP3 in chickens suppresses satellite cell differentiation^32^. Thus, it is likely that CSRP3 acts as a mechanosensor that plays a role in initiating resistance training-induced hypertrophy.

Adaptations to resistance training were also highly sex-specific. This difference in training response between sexes was mainly driven by substantial remodeling of intermediate filaments in females. Interestingly, intermediate filament proteins were also more abundant in untrained female skeletal muscle, indicating a greater reliance on intermediate filaments for muscle function in females. Disruption of intermediate filaments via mutation or knockout of desmin induces muscle weakness, disruption of myofibrils and myopathy^33,34^. In addition, desmin-null muscles also show less nuclei deformation upon mechanical stretch, which leads to absence of stretch-induced signaling^35^. Here, we are the first to identify sex-specific abundance and remodeling of intermediate filaments in skeletal muscle.

When trying to untangle the diverse differences between sexes and fiber types that may explain differences in training responsiveness our data suggest a role of translational capacity in mediating muscle fiber hypertrophy. As proteins involved in translation differed both between sexes and fiber types, this may be a potential mechanism of preferential type II fiber hypertrophy and limited resistance training-induced hypertrophy of type I fibers in females^9,36^. Likewise, greater translational capacity may also explain preferential type II fiber proteome remodeling in response to resistance training. This is likely due to protein synthesis being highly regulated at the translational level^37-39^. Therefore, the abundance of ribosomal proteins and eukaryotic initiation/elongation factors could be a limiting factor for resistance training-induced protein synthesis.

In summary, our proteomics workflow allowed mapping of the first fiber type-specific skeletal muscle proteome in females, and revealed remarkable differences to the male skeletal muscle proteome that were fiber type-specific. Likewise, resistance training-induced remodeling of the skeletal muscle proteome was also highly affected by sex- and fiber type. These results provide a resource to further investigate mechanisms of skeletal muscle adaptations to resistance training and underline the importance of assessing muscle adaptations in a sex- and fiber type-specific manner. As such, future research in the field of muscle physiology and muscle adaptations should strive to address research in a sex-specific manner, as research in males may not always be applicable to females.

## Methods

### Subjects

Twenty-four subjects (12 males and 12 females) completed the study. Inclusion criteria were between 18-45 years old, and body mass index (BMI) < 27. Before inclusion, each subject provided their written and oral informed consent. Exclusion criteria were smoking, chronic disease, use of prescription medication, and resistance training more than once per week in the 12 months leading up to the intervention. Subject characteristics are described elsewhere^9^. The study was approved by the local ethics committee of Copenhagen, Denmark (H-18007889) and registered on ClinicalTrials.gov (NCT04136457). All experimental procedures involving human subjects were performed at the Department of Nutrition, Exercise, and Sports, University of Copenhagen. The study complied with the 2013 declaration of Helsinki.

### Experimental protocol

The study consisted of an 8-week intervention, where subjects performed resistance training three times a week on non-consecutive days. The resistance training protocol is described elsewhere^9^. Before and three days after the intervention subjects attended an experimental day where leg lean mass was assessed by dual-energy X-ray absorptiometry (DXA) (Lunar iDXA, GE Healthcare, GE Medical systems, Belgium), a muscle biopsy was sampled from m. vastus lateralis, and one-repetition maximum (1RM) leg press was assessed. Subjects were instructed to replicate their dietary intake in the 24 h leading up to the experimental days. All experimental procedures were conducted at the same time of the day for each subject.

### Dual-energy X-ray absorptiometry

Two consecutive DXA scans were performed to account for scan-to-scan variation^40^. Prior to DXA scans subjects rested for 10 min in a supine position to ensure even distribution of body fluids^41,42^. Subjects were placed in the scanner undressed and euhydrated in a standardized supine position. The scanner was calibrated before scans using daily calibration procedures (Lunar ‘System Quality Assurance’).

### One-repetition maximum (1RM)

Prior to assessment of 1RM leg press a warm-up consisting of 6, 4 and 2 repetitions at 50, 70 and 80% of the anticipated 1RM was performed. For 1RM determination, Subjects then performed sets of one repetition of increasing weight, until the weight could not be lifted through the full range of motion. Three min of rest were provided between each successive attempt. The highest load successfully lifted was defined as 1RM. All 1RM determinations were made within five attempts.

### Muscle biopsies

Muscle biopsies were obtained under local anesthesia (lidocaine without epinephrine, Xylocaine 20 mg/mL, AstraZeneca) using a 4 mm Bergström needle (Stille, Stockholm, Sweden) with suction^43^. Immediately after collection, muscle biopsies were washed in sterile saline solution (0.9% NaCl) and quickly dissected free of visible blood, connective tissue, and fat. Thereafter, muscle biopsies were snap-frozen in liquid nitrogen, and stored at –80°.

### Preparation of muscle fiber pools

The frozen muscle biopsies were freeze dried for 48 h before dissection of single muscle fibers using fine forceps under a stereomicroscope in a temperature- and humidity-controlled room (humidity of ∼25%). A total of 40 fibers with a minimum length of 1 mm (1.5 ± 0.4 mm; mean ± SD) were dissected from each biopsy. Each fiber was dissolved in 15 µl of lysis buffer (1% SDC, 50 mM Tris-HCL pH 8.5, and ddH_2_O) and boiled for 10 min at 95° in a thermomixer. Thereafter, samples were sonicated using a bioruptor twin (Diagenode) for 10 min. To ensure fibers were at the bottom of the tube, samples underwent centrifugation (20.000 g, 1 min) between each step. A small fraction of each fiber was used to identify MYH expression by dot blotting with specific antibodies against MYH2 (mouse igM, 1:200, A4.1519, DSHB) and MYH7 (mouse igM, 1:200, A4.840, DSHB). For dot blotting, 1 ul (1/15 of a fiber lysate) was spotted onto two identical PVDF membranes. The membranes were blocked in 5% skimmed milk diluted in Tris-buffered saline with 0.1% Tween-20 (TBST) for 15 minutes, followed by incubation in primary antibody for 2 h at room temperature. One membrane was incubated in primary antibody against MYH2 and the other membrane was incubated in primary antibody against MYH7. Thereafter, membranes were incubated in horseradish peroxidase-conjugated secondary antibody (goat anti-mouse igG, 1:20.000; Bio-Rad) for 1 h at room temperature.

Primary antibodies were diluted in 3% BSA in TBST and secondary antibodies were diluted in 2% skimmed milk in TBST. Membranes were washed for 3 × 5 min between each step in the protocol. Dots were visualized using chemiluminescence (Immobilon Forte, Millipore) in a ChemiDoc XRS+ imaging system (Bio-Rad, CA, USA). The remaining sample of each fiber lysate was pooled according to MYH expression, resulting in pools of type I fibers that expressed only MYH7 and pools of type II fibers that expressed only MYH2. Hybrid fibers that expressed both MYH isoforms were excluded.

### Preparation of samples for proteomics analysis

Following determination of protein concentration via BCA protein assay (Thermo Fisher Scientific), 5µg of protein from each fiber type-specific lysate underwent reduction with DTT (final concentration of 10 mM) and alkylation with CAA (final concentration of 50 mM). Then, proteins were digested overnight using trypsin (Promega) and LysC (Wako) with an enzyme to protein ratios of 1:100 and 1:500, respectively. The following day, tryptic peptides were desalted using in-house crafted styrenedivinylbenzene reverse-phase sulfonate (SDB-RPS) StageTip columns and 200 ng of the resulting desalted peptides were loaded in Evotips according to the manufacturer’s instructions.

### Liquid chromatography tandem mass spectrometry

Proteomics measurements were conducted using LC-MS instrumentation comprising an Evosep One HPLC system^44^ (Evosep) coupled via electrospray ionization to a timsTOF mass spectrometer (Bruker). Peptides underwent separation on an 8 cm, 150 μM ID column packed with C18 beads (1.5 μm) (Evosep). The ‘60 samples per day’ chromatographic method was used to achieve peptide separation prior electrospray ionization through a CaptiveSpray ion source and a 10 μm emitter into the MS instrument. Samples were measured using a previously described DIA-PASEF workflow^45^. Specifically, the DIA-PASEF scan range was set at 400-1000 (m/z), the TIMS mobility range to 0.64-1.37 (V cm−2), and ramp and accumulation times were both set to 100 ms. To accommodate the ‘60 samples per day method’, A short-gradient method was used, which included 8 DIA-PASEF scans with three 25 Da windows per ramp, resulting in an estimated cycle time of 0.95 sec.

Raw spectra were analyzed by the search engine DIA-NN (version 1.878) using a library-based approach against an in-house muscle fiber-specific MS library containing 5000 proteins. Proteotypic peptides were used for protein group quantification, with the neural network set to double-pass mode. “Robust LC (high accuracy)” was chosen as quantification strategy as well as enabling the match between runs option, while the rest of parameters remained as default, including peptide length ranging 7-30 amino acids and precursor FDR set to 1%.

### Bioinformatics

Bioinformatics were conducted within the R environment (R version 4.3.2). Data were filtered such that only proteins that were quantified in 70% of the samples in at least one group (male type I, male type II, female type I, and female type II) were included for analysis. During filtering, samples from one female subject were excluded due to low protein abundance suggesting sample loss during sample preparation. Thereafter, data were log2 transformed and normalized by median scaling using the PhosR package. Categorical annotations for each protein were supplied in the form of GO terms of biological process (BP), cellular component (CC) and molecular function (MF), as well as UniProt Keywords. All annotations were extracted from the UniProt database. Principle component analysis was performed using the ppca method from the pcaMethods package. Differential expression analysis was performed using the limma package, with a false discovery rate (FDR) set to 5% using the qvalue package. Enrichment analysis of GO:BP terms was performed with the clusterProfiler package using a ranked protein list, with proteins ranked based on log2fold change obtained from differential expression analysis or pearson’s r from the correlation with change in fiber cross-sectional area (CSA)^9^. A FDR of 5% was chosen for enrichment analysis.

## Acknowledgments

We acknowledge Shah Mir Ahmed Bibi for his contribution to optimizing the dot blotting protocol. Mass spectrometry analyses were performed by the Proteomics Research Infrastructure (PRI) at the University of Copenhagen (UCPH), supported by the Novo Nordisk Foundation (NNF) (grant agreement number NNF19SA0059305).

## Author contributions

Conception and design: L.M, R.MJ, A.S.D, M.H. Performed experiments: L.M, R.MJ, J.S, S.J. Data analysis: L.M, R.MJ, S.J, B.S. Interpretation of results: All authors, Drafted the manuscript: L.M, Edited and revised the manuscript: All authors. All authors read and approved the final version of the manuscript.

## Supplementary material

**Table S1.** Proteomics data used for bioinformatics analysis

**Table S2.** Results of differential expression analysis between type I and type II fibers in males and females

**Table S3.** Results of main effect of sex from differential expression analysis between males and females

**Table S4.** Results of differential expression analysis between males and females in type I and type II fibers

**Table S5.** Results of main effect training from differential expression analysis between pre and post samples

**Table S6.** Results of differential expression analysis between pre and post samples in type I and type II fibers in males and females

## References

1 García-Hermoso, A. et al. Muscular strength as a predictor of all-cause mortality in an apparently healthy population: a systematic review and meta-analysis of data from approximately 2 million men and women. Arch Phys Med Rehabil 99, 2100–2113.e2105 (2018).

2 McLeod, J. C., Stokes, T. & Phillips, S. M. Resistance exercise training as a primary countermeasure to age-related chronic disease. Front Physiol 10, 645 (2019).

3 Phillips, S. M. & Winett, R. A. Uncomplicated resistance training and health-related outcomes: evidence for a public health mandate. Curr Sports Med Rep 9, 208–213 (2010).

4 Westcott, W. L. Resistance training is medicine: effects of strength training on health. Curr Sports Med Rep 11, 209–216 (2012).

5 Needham, D. M. Red and white muscle. Physiol Rev 6, 1–27 (1926).

6 Häkkinen, K., Komi, P. V. & Tesch, P. A. Effect of combined concentric and eccentric strength training and detraining on force-time, muscle fiber and metabolic characteristics of leg extensor muscles. Scand J Med Sci Sports 3, 50–58 (1981).

7 Staron, R. S. et al. Muscle hypertrophy and fast fiber type conversions in heavy resistance-trained women. Eur J Appl Physiol Occup Physiol 60, 71–79 (1990).

8 Tesch, P. A. Skeletal muscle adaptations consequent to long-term heavy resistance exercise. Med Sci Sports Exerc 20, S132–134 (1988).

9 Moesgaard, L., Jessen, S., Mackey, A. L. & Hostrup, M. Myonuclear addition is associated with sex-specific fiber hypertrophy and occurs in relation to fiber perimeter not cross-sectional area. J Appl Physiol (1985) 133, 732–741 (2022).

10 Beery, A. K. & Zucker, I. Sex bias in neuroscience and biomedical research. Neurosci Biobehav Rev 35, 565–572 (2011).

11 Koopman, R., Zorenc, A. H., Gransier, R. J., Cameron-Smith, D. & van Loon, L. J. Increase in S6K1 phosphorylation in human skeletal muscle following resistance exercise occurs mainly in type II muscle fibers. Am J Physiol Endocrinol Metab 290, E1245–1252 (2006).

12 Edman, S., Söderlund, K., Moberg, M., Apró, W. & Blomstrand, E. mTORC1 signaling in individual human muscle fibers following resistance exercise in combination with intake of essential amino acids. Front Nutr 6, 96 (2019).

13 Deane, C. S. et al. Proteomic features of skeletal muscle adaptation to resistance exercise training as a function of age. Geroscience 45, 1271–1287 (2023).

14 Roberts, M. D., et al. A novel deep proteomic approach in human skeletal muscle unveils distinct molecular signatures affected by aging and resistance training. bioRxiv (2023).

15 Robinson, M. M. et al. Enhanced protein translation underlies improved metabolic and physical adaptations to different exercise training modes in young and old humans. Cell Metab 25, 581–592 (2017).

16 Deshmukh, A. S. et al. Deep muscle-proteomic analysis of freeze-dried human muscle biopsies reveals fiber type-specific adaptations to exercise training. Nat Commun 12, 304 (2021).

17 Katsuma, Y., Swierenga, S. H., Marceau, N. & French, S. W. Connections of intermediate filaments with the nuclear lamina and the cell periphery. Biol Cell 59, 193–203 (1987).

18 Shah, S. B. et al. Structural and functional roles of desmin in mouse skeletal muscle during passive deformation. Biophysical Journal 86, 2993–3008 (2004).

19 Bellin, R. M., Huiatt, T. W., Critchley, D. R. & Robson, R. M. Synemin may function to directly link muscle cell intermediate filaments to both myofibrillar Z-lines and costameres. J Biol Chem 276, 32330–32337 (2001).

20 O’Neill, A. et al. Sarcolemmal organization in skeletal muscle lacking desmin: evidence for cytokeratins associated with the membrane skeleton at costameres. Mol Biol Cell 13, 2347–2359 (2002).

21 Rath, S. et al. MitoCarta3.0: an updated mitochondrial proteome now with sub-organelle localization and pathway annotations. Nucleic Acids Res 49, D1541–D1547 (2020).

22 Wackerhage, H., Schoenfeld, B. J., Hamilton, D. L., Lehti, M. & Hulmi, J. J. Stimuli and sensors that initiate skeletal muscle hypertrophy following resistance exercise. J Appl Physiol (1985) 126, 30–43 (2019).

23 Bernárdez-Vázquez, R., Raya-González, J., Castillo, D. & Beato, M. Resistance training variables for optimization of muscle hypertrophy: an umbrella review. Front Sports Act Living 4, 949021 (2022).

24 Kim, G. & Kim, J. H. Impact of skeletal muscle mass on metabolic health. Endocrinol Metab (Seoul) 35, 1–6 (2020).

25 Diaz-Canestro, C., Pentz, B., Sehgal, A. & Montero, D. Sex differences in cardiorespiratory fitness are explained by blood volume and oxygen carrying capacity. Cardiovasc Res 118, 334–343 (2021).

26 Murphy, W. G. The sex difference in haemoglobin levels in adults - mechanisms, causes, and consequences. Blood Rev 28, 41–47 (2014).

27 Toivola, D. M., Tao, G. Z., Habtezion, A., Liao, J. & Omary, M. B. Cellular integrity plus: organelle-related and protein-targeting functions of intermediate filaments. Trends Cell Biol 15, 608–617 (2005).

28 Cornelison, D. D. & Wold, B. J. Single-cell analysis of regulatory gene expression in quiescent and activated mouse skeletal muscle satellite cells. Dev Biol 191, 270–283 (1997).

29 Boateng, S. Y., Senyo, S. E., Qi, L., Goldspink, P. H. & Russell, B. Myocyte remodeling in response to hypertrophic stimuli requires nucleocytoplasmic shuttling of muscle LIM protein. J Mol Cell Cardiol 47, 426–435 (2009).

30 Gunkel, S., Heineke, J., Hilfiker-Kleiner, D. & Knöll, R. MLP: a stress sensor goes nuclear. J Mol Cell Cardiol 47, 423–425 (2009).

31 Kong, Y., Flick, M. J., Kudla, A. J. & Konieczny, S. F. Muscle LIM protein promotes myogenesis by enhancing the activity of MyoD. Mol Cell Biol 17, 4750–4760 (1997).

32 Han, S. et al. Knockdown of CSRP3 inhibits differentiation of chicken satellite cells by promoting TGF-β/Smad3 signaling. Gene 707, 36–43 (2019).

33 Li, Z. et al. Desmin is essential for the tensile strength and integrity of myofibrils but not for myogenic commitment, differentiation, and fusion of skeletal muscle. J Cell Biol 139, 129–144 (1997).

34 van Spaendonck-Zwarts, K. Y. et al. Desmin-related myopathy. Clin Genet 80, 354–366 (2011).

35 Palmisano, M. G. et al. Skeletal muscle intermediate filaments form a stress-transmitting and stress-signaling network. J Cell Sci 128, 219–224 (2015).

36 Abou Sawan, S., et al. Satellite cell and myonuclear accretion is related to training-induced skeletal muscle fiber hypertrophy in young males and females. J Appl Physiol (1985) 131, 871–880 (2021).

37 Jefferson, L. S. & Kimball, S. R. Translational control of protein synthesis: implications for understanding changes in skeletal muscle mass. Int J Sport Nutr Exerc Metab 11 **Suppl**, S143-149 (2001).

38 Baar, K. & Esser, K. Phosphorylation of p70(S6k) correlates with increased skeletal muscle mass following resistance exercise. Am J Physiol 276, C120–127 (1999).

39 Merrick, W. C. Mechanism and regulation of eukaryotic protein synthesis. Microbiol Rev 56, 291–315 (1992).

40 Zemski, A. J. et al. Same-day vs consecutive-day precision error of dual-energy X-ray absorptiometry for interpreting body composition change in resistance-trained athletes. J Clin Densitom 22, 104–114 (2019).

41 Berg, H. E., Tedner, B. & Tesch, P. A. Changes in lower limb muscle cross-sectional area and tissue fluid volume after transition from standing to supine. Acta Physiol Scand 148, 379–385 (1993).

42 Cerniglia, L. M., Delmonico, M. J., Lindle, R., Hurley, B. F. & Rogers, M. A. Effects of acute supine rest on mid-thigh cross-sectional area as measured by computed tomography. Clin Physiol Funct Imaging 27, 249–253 (2007).

43 Bergstrom, J. Percutaneous needle biopsy of skeletal muscle in physiological and clinical research. Scand J Clin Lab Invest 35, 609–616 (1975).

44 Bache, N. et al. A novel LC System embeds analytes in pre-formed gradients for rapid, ultra-robust proteomics. Mol Cell Proteomics 17, 2284–2296 (2018).

45 Meier, F. et al. diaPASEF: parallel accumulation-serial fragmentation combined with data-independent acquisition. Nat Methods 17, 1229–1236 (2020).

